# GMMchi: Gene Expression Clustering Using Gaussian Mixture Modeling

**DOI:** 10.1101/2022.02.14.480329

**Authors:** Ta-Chun Liu, Peter N. Kalugin, Jennifer L. Wilding, Walter F. Bodmer

**Affiliations:** Cancer and Immunogenetics Laboratory, Weatherall Institute of Molecular Medicine, Department of Oncology, University of Oxford, John Radcliffe Hospital, Oxford OX3 9DS, United Kingdom; Harvard Medical School, Boston, MA 02115, USA

## Abstract

**Motivation:** Cancer evolution consists of a stepwise acquisition of genetic and epigenetic changes, which alter the gene expression profiles of cells in a particular tissue and result in phenotypic alterations acted upon by natural selection. The recurrent appearance of specific genetic lesions across individual cancers and cancer types suggests the existence of certain “driver mutations,” which likely make up the major contribution to tumors’ selective advantages over surrounding normal tissue and as such are responsible for the most consequential aspects of the cancer cells’ gene expression patterns and phenotypes. We hypothesize that such mutations are likely to cluster with specific dichotomous shifts in the expression of the genes they most closely control, and propose **GMMchi**, a Python package that leverages Gaussian Mixture Modeling to detect and characterize bimodal gene expression patterns across cancer samples, as a tool to analyze such correlations using 2x2 contingency table statistics.

**Results:** We confirm that **GMMchi** robustly and reliably extracts bimodal patterns from both colorectal cancer (CRC) cell line-derived microarray and tumor-derived RNA-Seq data and verify previously reported gene expression correlates of some well-characterized CRC phenotypes. Using well-defined simulated data, we were able to confirm the robust performance of GMMchi, reaching 85% accuracy with a sample size of n = 90. We were also able to demonstrate a few examples of the application of **GMMchi** with respect to its capacity to characterize background florescent signal in microarray data, filter out uninformative background probe sets, as well as uncover novel genetic interrelationships and tumor characteristics. Our approach to analysing gene expression analysis in cancers provides an additional lens to supplement traditional continuous-valued statistical analysis by maximizing the information that can be gathered from bulk gene expression data.

**Availability:** The Python package **GMMchi** and our cell line microarray data used in this paper is available for downloading on GitHub at https://github.com/jeffliu6068/GMMchi

## 1 INTRODUCTION

Gene expression data are one of the most widely used tools in multiple aspects of understanding cell biology. There is a range of tools developed to evaluate expression levels including the microarray and RNA-seq technologies. Gene expression data produced across all platforms inherently results in a continuous-valued dataset representing the levels of expression of individual genes. The field of computational gene expression analysis is thus an analysis of continuous distributions and the correlative understanding of between genes or sets of genes activities within a biological context.

In cancer biology, tumorigenesis is driven by stable or meta-stable alterations at the genetic or epigenetic levels. Many of these changes probably lead to minor changes in the phenotype of a cell and minimal, if any, reproductive advantage. Some genetic mutations or meta-stable epigenetic expression changes that are common across different types of cancers do however lead to phenotypes that confer considerable reproductive advantages characteristic of a tumor, such as uncontrolled proliferation countering the control of normal differentiation, anti-apoptosis, and immune evasion (Bailey et al. 2018). These mutations and gene expression changes causing such strong reproductively advantageous phenotypic changes are known as driver mutations or epigenetic drivers because it is the successive accumulation of these advantageous genetic or epigenetic changes that drives the major development and progression of a cancer.

A single selectively advantageous mutation or epigenetic change may often lead to changes in the expression level of a whole set of other genes. This will give rise to a cluster of gene expression changes associated with such a particular mutation or meta-stable epigenetic change. The expectation is then that each expression distribution of a gene in such a cluster, over a range of cancers, will be bimodal with the mutation or epigenetic change associated with one mode and the corresponding normal version with the other mode of the bimodal distribution (Wang et al 2009).

The more closely a change in a gene expression level is associated with a given mutation or epigenetic change, the more likely it is to give rise to a bimodal distribution differentiating wild-type and mutation. More generally, on the other hand, the more striking is the distinction between two particular phenotypes over a range of cancers, the more likely it is to have a measurable effect on the biology of the cell, and the more likely it is to be due to a single mutation or stable epigenetic change. This is simply an extension of the principle by which Mendel established his laws of inheritance. Thus, the more distinctive is the bimodality of the expression of a gene over a range of cancers, the more likely it is to be due to, or strongly associated with, a genetic mutation or meta-stable epigenetic change that is giving an advantage to the outgrowth of the tumour, namely a cancer driver. The interest is then to identify striking bimodal gene expression distributions, over a given range of cancers, that lead back to novel mutations or epigenetic drivers. Here this is approached by fitting mixtures of two normal distributions, called Gaussian Mixture Models or GMM, to the gene expression data.

As shown in Figure 1, the most common distribution patterns found in gene expression data are unimodal or bimodal distributions consisting, respectively, of 1 or 2 normal (Gaussian) distributions. The distributions shown in Fig1 are histograms of gene expression data over a collection of 78 cell lines derived from different colorectal cancers (CRC), with the number of cancer cancers on the y-axis and the expression levels in log_2_ from micro-array data on the x axis. The distribution for the gene *GJC2* (coding for a gap junction protein) is clearly unimodal and well fitted by a single normal distribution, while that for gene *CDX1* (coding for a homeobox transcription factor) is clearly well fitted by a mixture of two normal distributions. One of the major challenges for GMM is to test the validity of the assumption that the data either fit a unimodal normal distribution or a mixture of two normal distributions. In Figure 1 the distribution for the gene *CDH1*, encoding E-cadherin, only fits a normal distribution at its upper end, while having a ‘tail’ of lower values that do not fit a normal distribution. In this paper, we describe a systematic approach to a novel GMM method capable of identifying normal and non-normal components across data on samples of any given form of expression data and give examples of the method’s application to expression data on a panel of CRC derived cell lines.

**Figure 1.**
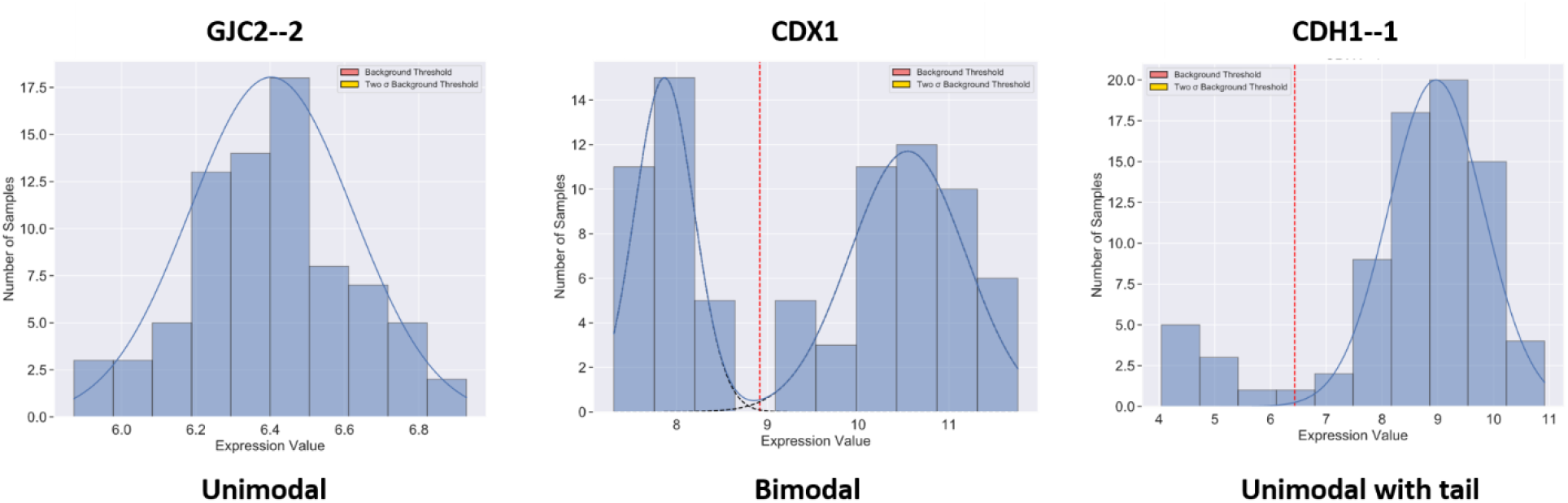
Common Distributions in Gene Expression Analysis: Examples of common histogram distributions seen in gene expression data from a panel of 78 colorectal cancer (CRC) derived cell lines. Expression levels based on microarray analysis are given as log_2_ on the x-axis, and numbers of samples with given expression levels on the y-axis. The continuous curves are fitted normal distributions and the vertical dotted red line marks the best estimate for separating low and high levels of expression, derived as described later. The mRNA for the gene coding GJC2, a gap junction protein, exhibits a unimodal distribution representing a single Gaussian distribution while the mRNA for the gene coding CDX1, a homeobox protein, exhibits a bimodal distribution representing two distinct Gaussian distributions. The gene for CDH1, encoding E-cadherin, shows a unimodal distribution with a tail of low expression values, whose estimation is one of the main challenges for GMM.

## 2 Pipeline development: introduction

### 2.1 Gene Expression Categorization

The primary objectives of this analytical approach are twofold:

1. Develop an unsupervised method for determining whether a gene’s expression pattern across a set of cell or tissue samples is unimodal or bimodal. If the expression data are a mixture of normal and non-normal components, separate the samples into normal and non-normal components before making the appropriate cutoff separating the samples into individual subgroups.
2. For genes with bimodal expression patterns, analyze their patterns of association into sub-group clusters using 2x2 contingency table analysis and explore the associations of these clusters with gene mutations and biological properties, such as patterns of differentiation.

This analytical pipeline takes standard normalized and batch-removed gene expression datasets as input obtained either from microarrays or by RNA-Seq.

### 2.2 Gaussian Mixture Modeling

Gaussian Mixture Modeling (GMM) is performed using the **sklearn.mixture** package in Python. The expression pattern of each gene or probe set is examined as a univariate distribution across the set of samples in the dataset. One or two Gaussian distributions are fitted to each distribution, in the latter case allowing for unequal sub-distribution fractions and variances, using the expectation-maximization algorithm built into **sklearn.mixture.GMM**. For bimodal gene expression distributions, each sample is given a value 1 or 2 according to the sub- distribution it more likely belongs to, based on the posterior probability outputted by GMM.

Model selection, in particular the choice of either one- or two-Gaussian fitted distributions, is based on maximization of the Bayesian Information Criterion (BIC) defined in Equations (1) or, its approximation, (2):

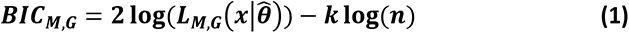

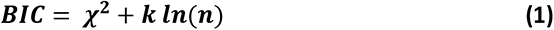

where 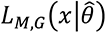 is the maximized likelihood function of model *M* with *G* components, with maximizing parameters 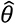, determined through the EM algorithm, *n* is the sample size, and *k* is the number of estimated parameters (Dempster *et al*., 1977). The BIC is a log-likelihood function, with a penalty term that increases with the number of parameters to counteract overfitting, and which has been shown to perform well in a range of applications (Schwarz, 1978; Fraley and Raftery, 1998). BIC is calculated for deciding between fitting our data with one or two Gaussian components. The model with the lower BIC value is selected.

### 2.3 Non-normally Distributed Tail Problem

The challenges faced with directly applying GMM to expression data analysis and the identification of bimodal distributions lie within the nature of gene expression datasets. Although expression data like many other datasets mostly behave as Gaussian distributions after log_2_ transformation (Quackenbush 2002), a large proportion of gene expression data exhibits a pattern where there is a, sometimes quite long, tail of outliers outside the normal distributions which contain most of the data, as exemplified in Figure 1. We call this the ‘tail problem’. This non-normally distributed tail is inadequately dealt with by GMM which assumes input data are all expected to be mixtures of normal distributions.

An example of the tail problem in the returned output of GMM is illustrated in Figure 2.

**Figure 2.**
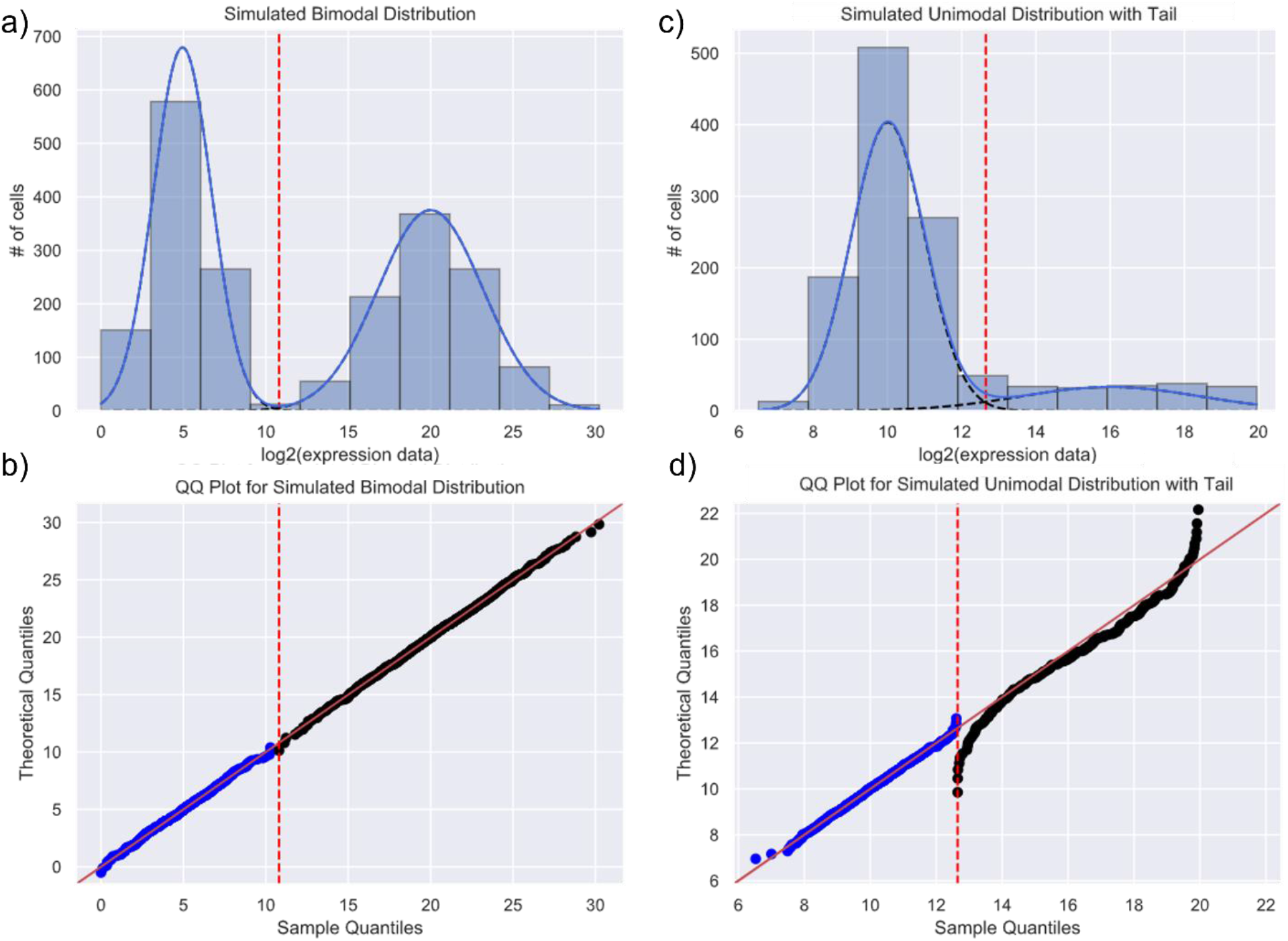
GMM on a Simulated Mixture of Normal Distributions: (a) shows a mixture of two. Gaussian components fitted to a simulated bimodal distribution using sklearn.mixture. The red dotted line represents the separation between the two fitted normal components. In the case of two normal distributions, the red dotted line is the intersection between the two distributions where the probability of a data point categorized under either of the two distributions crosses over from to another. (b) is a Q-Q plot that measures the normality of the fitted data by plotting the theoretical quantile of a normal distribution with the computed mean and variance against the sample quantile of the input data. Normality is visualized by the scatterplot lying on the 45- degree line suggesting the quantiles of the theoretical and sample data behave similarly. (a) and (b) are examples of a bimodal distribution adequately identified by GMM. (c) shows a simulated bimodal distribution with a ‘tail’, representing a non-normal spread of outliers inadequately fitted by GMM. Compared to (b) where the dots lie on the 45-degree line, (d) shows how a non-normal tail can skew the alignment of dots on the 45-degree line of the Q-Q plot. The slight uptick of blue dots near the red line represents the small overlap between the tail on the right and the normal distribution on the left where the assignment to the distribution to which each datapoint belongs is less well defined.

The dashed blue lines in Figures 2a and 2c are the GMM fitted normal distributions, while Figures 2b and 2d are the corresponding Q-Q plots, which show whether the quantiles of the fitted distributions are as expected for a normal distribution. In Figure 2b, the data lie perfectly on the 45^0^ diagonal lines when plotted against the theoretical normal distributions, indicating normality for both components of the sample distribution. On the other hand, in Figure 2d, the line for the righthand tail distribution is markedly skewed from the diagonal at both ends, strongly showing the non-normality of the tail distribution.

## 3 A GMM with a Chi-Square Pipeline: GMMchi

To deal with the tail problem we need to distinguish between a good GMM fit of two normal distributions with different variances and a poor over fit due to a mixture of one or two normal distributions with non-normal tail distributions. To make this distinction we add iterative Chi-square fitting to GMM and call this GMMchi. There are four steps to this iterative optimization process: dynamic binning, tail identification, iterative tail trimming, and a final condition criterion. We now address each step of the pipeline in turn.

### 3.1 Dynamic binning

The most important factors that determine how a series of individual data on a single continuous measurement are presented in a histogram are the number of bins and their widths. This is a particular problem when the number of individual observations is not very large. Both the bin width and the number of bins create problems for the iterative testing of the fit of the observed data to estimated normal distributions.

To create a fully automated pipeline, we have incorporated the Mann and Wald bin criterion (Bowman, D’Agostino, and Stephens 1988) defined in Equation (3):

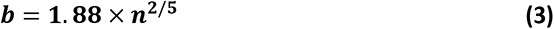

Here *b* is the number of bins while *n* is the number of samples. This method provides a consistent approach to deciding on the number of bins that best represents the data.

Next, since the iterative process within GMMchi is based on the chi-square goodness of fit test as the main criterion for measuring the fit of the mixed normal distribution model to the observed data, it is important for the validity of the Chi-square test to have at least 5 measurements in each bin. This is achieved by an algorithm called dynamic binning, which involves automatically combining bins while applying the least manipulation to the histogram for ensuring optimal results of the underlying chi-square test within GMMchi. The three-step process is as follows:

1. Use the Mann and Wald bin criterion to determine the number of bins *b*. Using *b* and the range of our data we can then propose that initially the width of our bins be 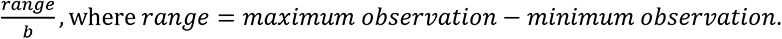
2. Quantify the number of observations per bin.
3. Starting from the upper and lower ends of the distribution, combine adjacent bins that have < 5 observations until all bins have at least 5 observations.

The results of dynamic binning are illustrated in Figure 3. As shown, the dynamic binning ensures there are more than 5 observations in every bin and so the validity of the chi-square goodness of fit test used in the subsequent steps. Dynamic binning is thus applied to all data before GMM fitting.

**Figure 3.**
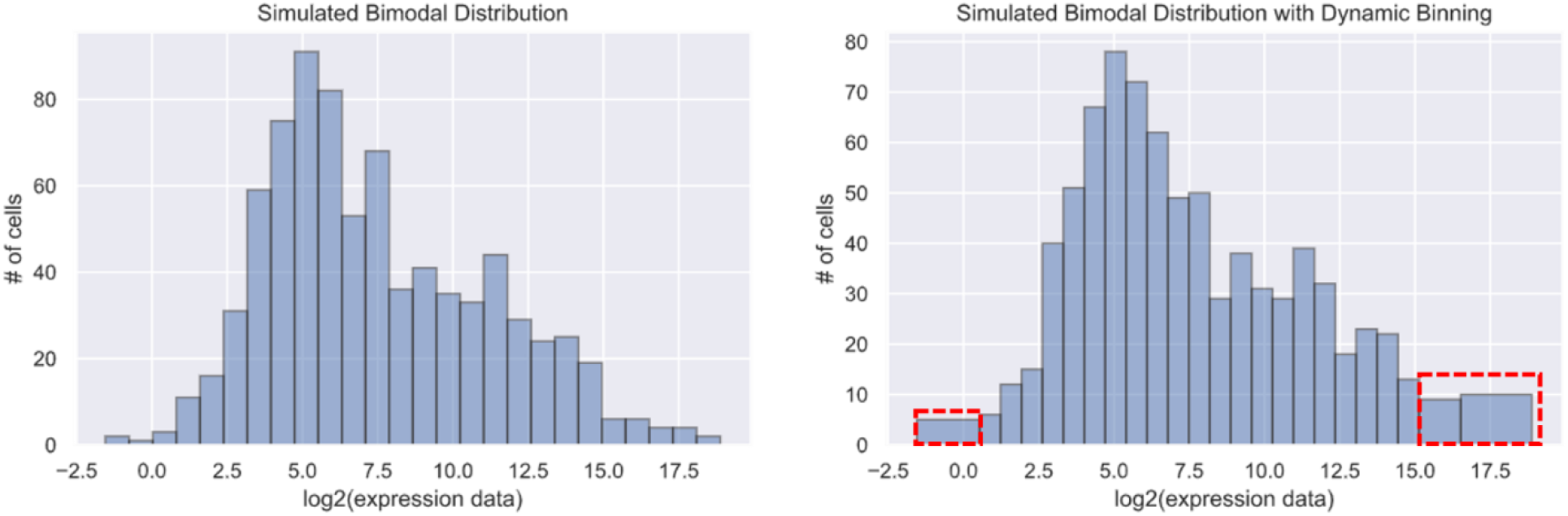
Dynamic Binning: Based on simulated data that creates a bimodal distribution. The mean and variance of the simulated 400 random samples in each histogram are 5, 10, and 3, 10, respectively. On the left is the histogram before applying dynamic binning. Samples within each bin in the far left and right of the left histogram are less than 5 thus falling below the prerequisite of applying the chi-square goodness of fit test. On the right, dynamic binning solves this by dynamically combining bins without over-manipulating the overall distribution. In this example, only 15 samples were combined via dynamic binning. The bins combined via dynamic binning are boxed in red dashes.

### 3.2 Tail Criterion

The next step deals with the tail problem. Our pipeline first identifies whether the data are a mixture of normal distribution(s) or a mixture of non-normal tails and normal distribution(s). If the data fit the non-normal tail criterion, GMMchi will iteratively remove one data point at a time from the extreme end of the tail while fitting the remaining points with GMM. At each tail point removed, the χ^2^ value for the remaining points as well as the BIC will be calculated to determine the best fitting outcome. The χ^2^ value is defined in Equation (4):

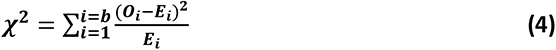

The χ^2^ value is thus the sum of the squared difference between the observed number, ***O_i_*** in the ***i^th^*** area under the curve, and ***E_i_***, the estimated number based on the currently fitted single or bimodal normal distributions, divided by ***E_i_***.

To determine whether there is a non-normally distributed tail the overall input data will first be fitted by GMM after applying dynamic binning. The returned output of the GMM consists of the number of normal components fitted to the data based on BIC, the posterior probability of each data point (x), and the characteristics of each component distribution, namely the mean (µ) and the variance (σ^2^). To determine at any given point in the pipeline whether there is overfitting of the model with a non-normal tail problem, we have devised a criterion based on the goodness of fit testing.

Tails have characteristic flat, spread-out distributions that are the outliers of the input data. Under GMM, tails will be fitted as normal distributions but will be expected to have poor fits to a normal distribution. Thus, the χ^2^ value from fitting 2 normal distributions to data without a tail will be lower, namely have a better fit, than fitting two normal distributions to data with a non-normal tail.

Based on this assumption, we can derive a tail criterion based on how well the model fits the data. We have selected the chi-square goodness of fit test as our test for model fitness due to its simplicity and generalizability. Compared to the BIC, chi-square goodness of fit is not directly affected by the total sample size involved in the penalizing term of the BIC, which changes with each iteration involved in determining the tail (see below), so that the computed chi-square values are comparable across different iterations of the pipeline. This is an important attribute that allows the comparison of the model fit across sequential iterations when selecting the best fit.

To determine the acceptable χ^2^ threshold value for tail detection, we plotted the distribution of the χ^2^ values of all our GMM bimodally fitted genes in the 78 -cell line panel, as shown in Figure 4. The degrees of freedom used to calculate the χ^2^ values for the estimated distribution are given in Equation (5):

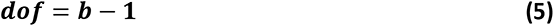

**Figure 4.**
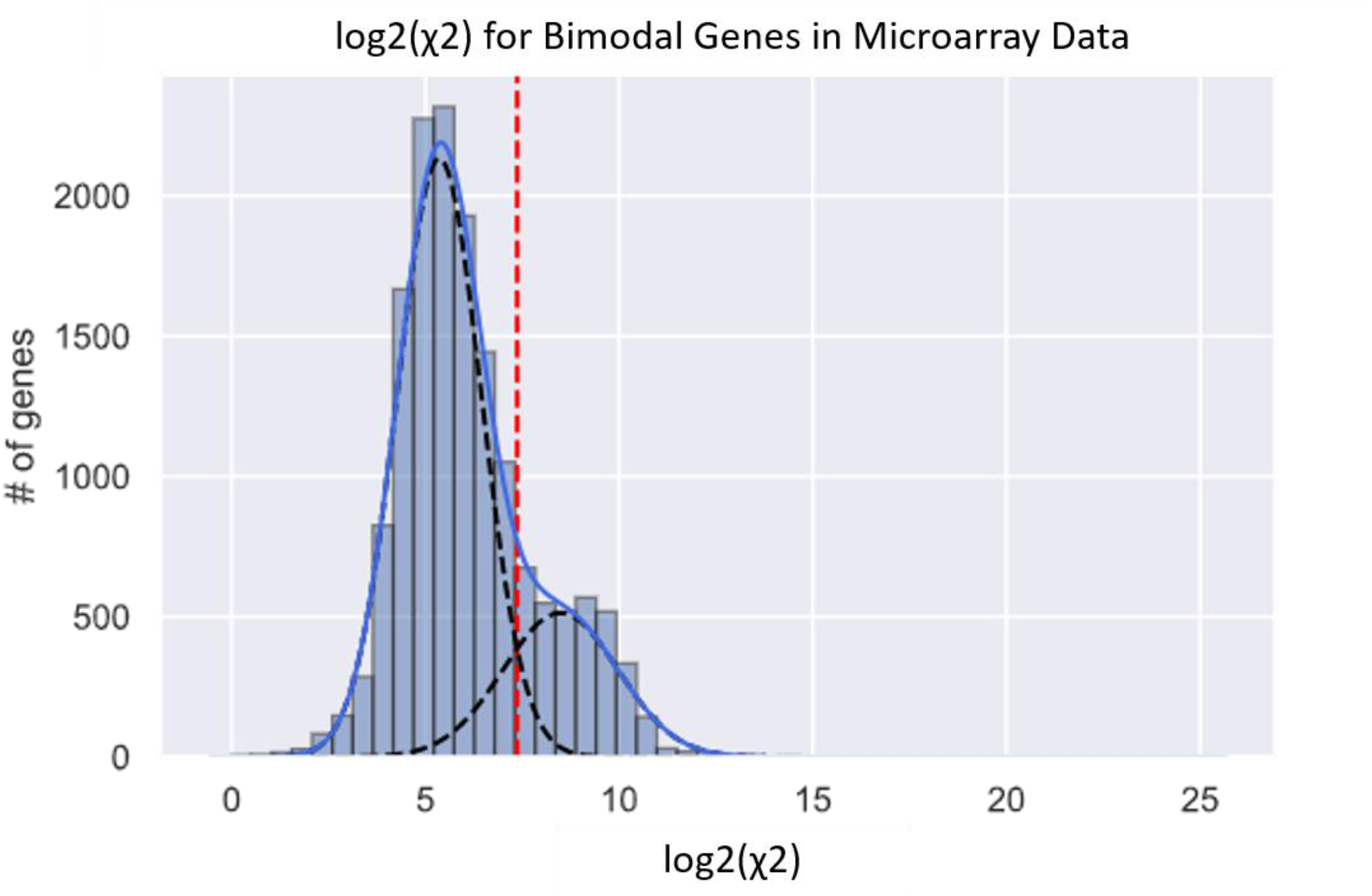
Determining the Chi-Square Threshold Value for Identifying a Poorly Fitted Tail: This histogram is the distribution of the spread of the log2-transformed χ^2^ values of all genes identified by GMM as bimodal in our panel of 78 cell lines. The histogram has a clear bimodal distribution with two normal components that are well fitted by GMM estimation. We assume the two distributions represent: 1) On the low side, the well-fitted distributions with lower χ^2^ values, and 2) the higher χ^2^ values indicating inadequately fitted distributions containing non-normal tail components. Using the estimated mean of the upper distribution of 8.48 and a standard deviation of 1.5, we estimate a lower threshold of 5.52 for the log_2_ χ^2^ values that indicate the presence of a non-normal tail at a false-negative rate of 5%.

Since ***dof*** are calculated based on the number of bins ***b***, they decrease very gradually as we remove 1 datapoint at each iteration because tails are outliers of the normal distribution and so only a limited number of datapoints are usually in the tail. This effect on the decrease of ***dof*** has, therefore, a minimal effect on the χ^2^ values.

The histogram of all χ^2^ values shows a clear bimodal distribution with two normal components fitted by GMM estimation (Figure 4). Assuming that the upper normal distribution represents genes with non-normal tails, hence a higher χ^2^ value, the lower normal distribution represents well-fitted distributions with lower χ^2^ value. Based on this result, we can reasonably assume any χ^2^ value larger than 2 ^5.52^ indicates the presence of a non-normal tail.

### 3.3 Iterative Tail Trimming (ITT)

After determining whether a distribution has a non-normal tail, our next goal is to identify the correct threshold that separates the tail from the main distribution. As discussed in the previous section, the tail criterion will label the result as ‘pass’ or ‘tail’. If the model is labeled as ‘pass’, the model passes the criterion and the result of the GMM will be respected and returned as the final output. If the model is labeled as a ‘tail’ we assume that the initial GMM fit to the data was inadequate and that the data consists of a mixture of normal distribution(s) with a non-normal tail that should be subjected to iterative tail trimming.

Iterative tail trimming (ITT) is an optimization process where the algorithm searches for the best fitting datapoint where the cutoff can be made to separate the tail from the rest of the normally distributed data. The process is as follows:

1. At each iteration, we begin by removing one data point from the tail, perform dynamic binning to adjust the appropriate number of observations per bin, and fit the resulting distribution with GMM.
2. Then, we determine the chi-square goodness of fit (χ^2^) using parameters from the output to determine how well the revised model fits the input data.
3. We next compare χ^2^ _n_ and χ^2^ _(n-1)_, the χ^2^ values of the current iteration (n) and the previous iteration (n-1), as well as the BIC score to determine whether the model is fitting better than the previous fit. We use the BIC score to determine whether a lower χ^2^ value is also the least complex model with the best fit.
4. The algorithm iterates 1), 2), and 3) until no other conditions produce a lower χ^2^ and BIC.

The result of ITT is thus a series of χ^2^ values representing the fit of the model at each iterative step. The lower the χ^2^ value the better the model fits. Even though BIC is a criterion that is equal to the χ^2^ value with a penalizing term, as shown in Equation (2), the reason why we are unable to depend solely on BIC is because the penalizing term is dependent on the sample size. Since we are removing 1 datapoint at each iteration during ITT, the sample size *n* is constantly decreasing making BIC a relative value that is only relevant when comparing inputs with the same sample size. This means that the χ^2^ value itself is an absolute number we use to compare across all data points to determine how well the new model fits and the BIC values are relative terms used to compare, within the same sample data, what is the best number of fitted distributions.

### 3.4 Multiple Conditions for Iterative Tail Trimming

Having established Iterative Tail Trimming (ITT), there is another important factor to consider before applying ITT. There are four basic possible distribution patterns a tail problem can give rise to in a GMMchi, as illustrated in Figure 5:

1. One normal distribution plus a low tail
2. One normal distribution plus a high tail
3. Two normal distributions plus an extreme low tail
4. Two normal distributions plus an extreme high tail

**Figure 5.**
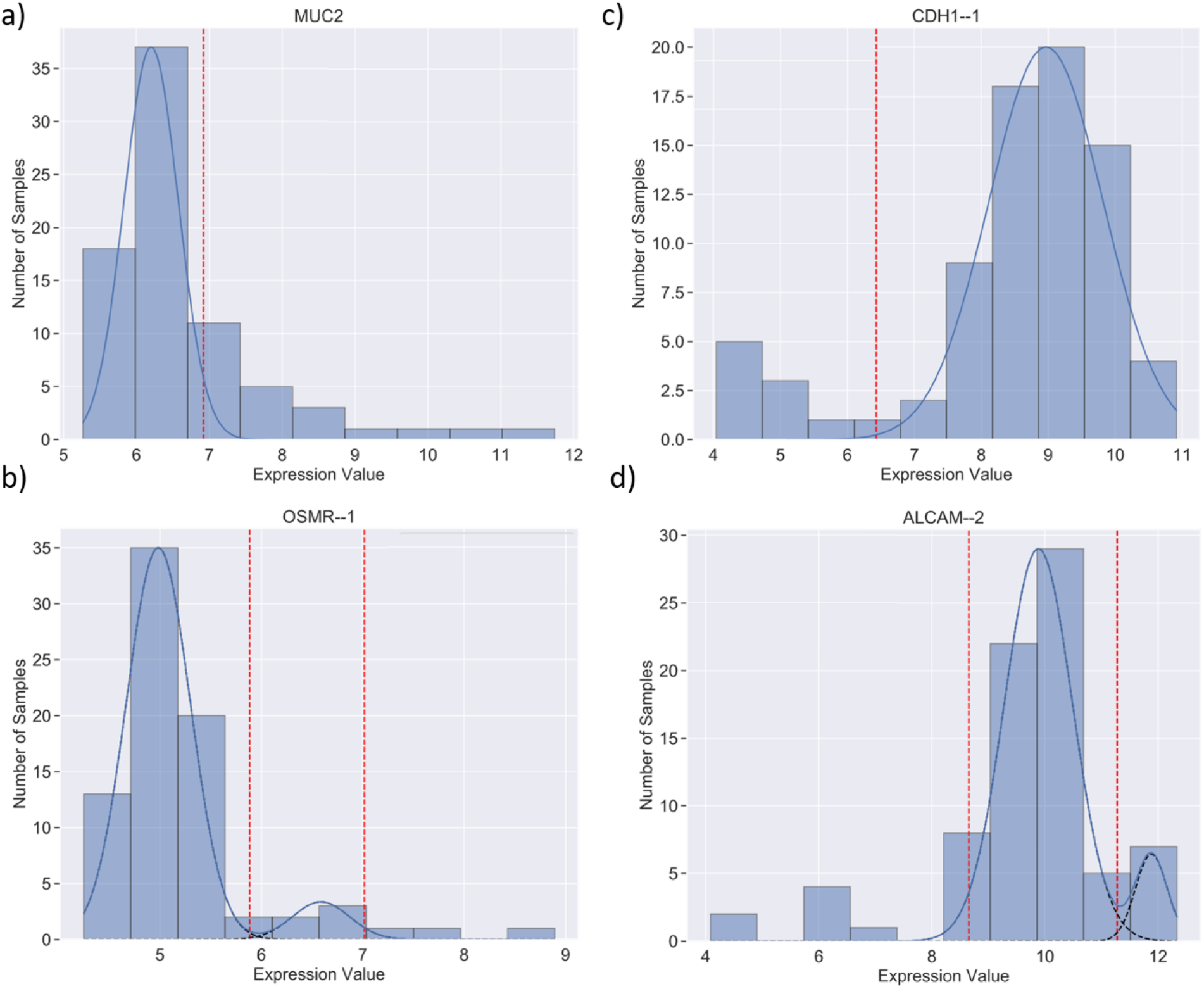
Examples of Different Distributions Found in Gene Expression Analysis: (a), (b), (c), and (d) demonstrate good examples of a unimodal distribution with a right tail, a bimodal distribution with a right tail, a unimodal distribution with a left tail, and a bimodal distribution with a left tail, respectively.

The algorithm takes all possible scenarios listed above into account in performing ITT on the data. This process runs ITT removing data from both low or high tails if the χ^2^ value is >2 ^5.52^ and fitting the remaining data with one or two mixtures of Gaussian distributions, covering all four possible scenarios mentioned above. The cutoff that determines where each tail starts is based on the intersection of the two overlapping normal distributions. This ensures all scenarios are well-considered within our model and the output represents the optimal tail trimming of the best fitting model. Possible scenarios including unimodal or bimodal distribution with both upper and lower tails are not considered in GMMchi due to their rare occurrence in gene expression data.

### 3.5 Final criterion for Determining the Cut-off that Separates the Main Distributions from the Tails: Returned Output of GMMchi

The final criterion is determined by identifying the lowest chi-square value, which indicates the best fit, and the lowest BIC score, which suggests the least complex model, across all 4 possible scenarios. This results in the definition of the tail data points and so the definition of the boundaries between the estimated normal distributions and the tails. The last step, when there is a well fitted bimodal distribution after omitting the tail(s), is to show the boundary between the high and low values of these two distributions by their point of intersection.

The resulting best fit can be either the original fit, indicating that a mixture, usually of two normal distributions, best represented the data, or a new fit consisting of a mixture of normal distributions and non-normally distributed tails.

The output of GMMchi includes the categorization of input data across all samples and a set of the means and variances that describe the fitted distributions.

The visual output consists of 4 graphs as in Figure 6. Figure 6a is the distribution of the input data (shown as a histogram) with the fitted model (shown as the line graph mapped onto the histogram). Figure 6c is a histogram of the percentage of each categorized group. Figure 6b is a graph of the BIC score that decides the number of components at each iterative fit and Figure 6d is a graph of the χ^2^ value that decides the best-fitting model at each iterative fit and eventually the best overall fitting model, which has the lowest χ^2^ value reached after removing 9 datapoints from the tail.

**Figure 6.**
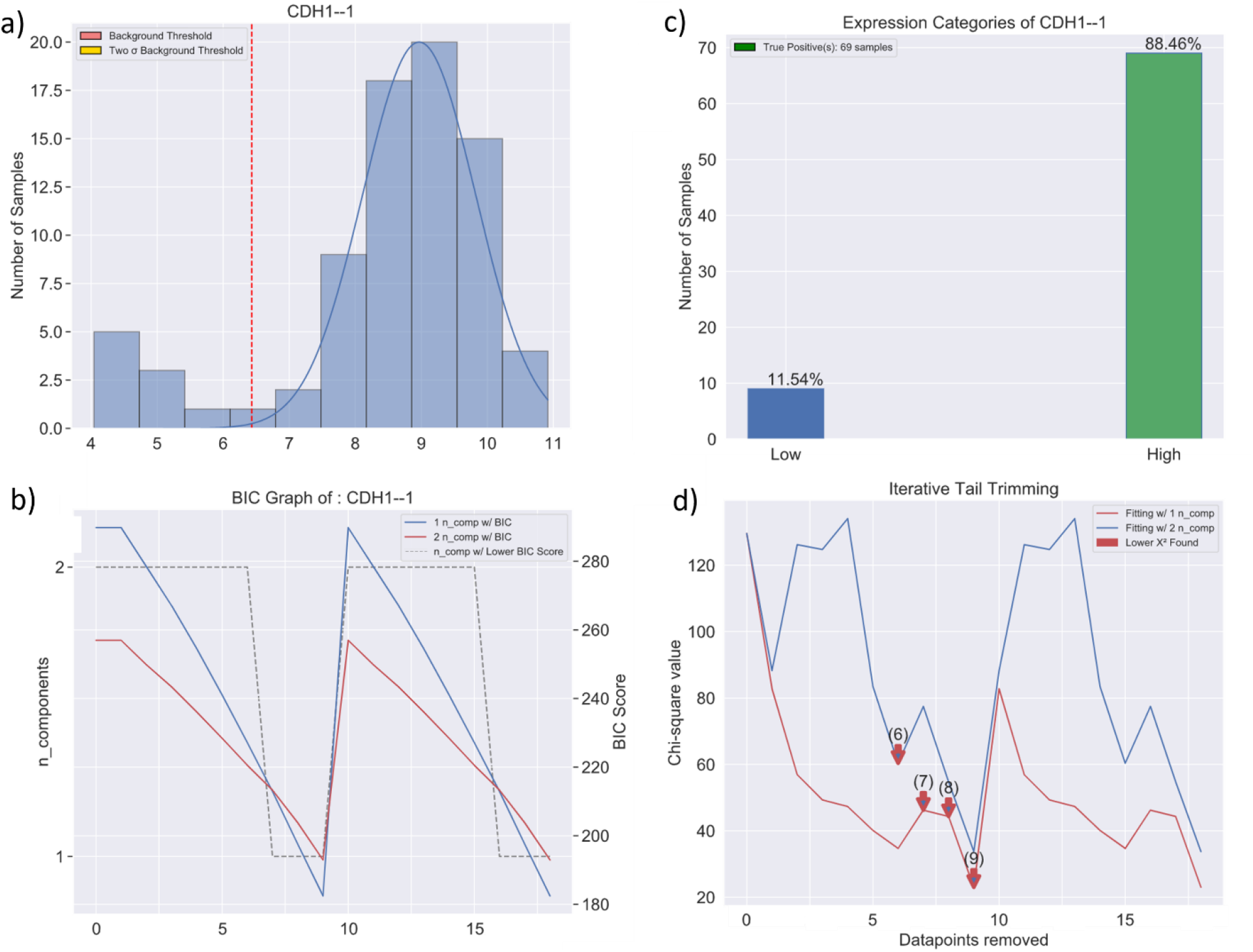
Example Output of the GMMchi Python Package: (a) is the log2-transformed distribution of the gene expression level of CDH1 (E-Cadherin) with its fitted normal components along with a red vertical cutoff from the tail. (b) is the BIC score. The red being the BIC score for fitting 2 normal components, the blue being the BIC score for fitting 1 normal component and the grey dashed line indicating which model to pick at each iterative step. (c) is a bar graph showing the percentage of samples under each categorized group. (d) is a graph showing the χ^2^ value calculated at each iterative step during iterative tail trimming. The red line represents the χ^2^ value of fitting 1 normal component, the blue line represents the χ^2^ value of fitting 2 normal components, and the red arrows indicate when a better fit is found. The best fit with the lowest χ^2^ value and the lowest BIC is with 9 low tail values removed and just one normal distribution for the remaining data.

## 4 GMMchi validation

To test the robustness and sensitivity of the GMMchi pipeline, we performed two separate simulations.

### 4.1 Simulation test 1

The purpose of this simulation is to assess how effective GMMchi is at identifying a simulated mixture of normal and non-normal tails with an increasing spread of the tail. The spread or size of the tail is defined as its range. There are two components to this simulation:

1. Unimodal case: Mixture of 1 normal distribution + 1 non-normal tail
2. Bimodal case: Mixture of 2 normal distributions + 1 non-normal tail

We randomly generate normal distributions choosing means at random between 5 and 10 and standard deviations at random between 0.5 and 5. The total number of data points is chosen at random to be between 50 and 150. The number of data points in the tail is randomly chosen to be between 5-25% of the total number of data points. To test how well GMMchi performs with an increasing spread of the tail, we choose the lower bound of the tail to be *lower bound* = *m* + 2 × *std* where *m* and *std* are respectively the mean and standard deviation of the normal component. This ensures that the tail starts at the upper end of the distribution when there is just one normal distribution. For 2 normal distributions, we use the *m* and *std* of the upper distribution and start the tail at the upper end of this second distribution. The upper bound is determined by *upper bound* = *lower bound* + *n × std* where *n* is increased incrementally from 1 to 12 to give a larger spread of the tail with each set of simulations as *n* increases. We then generate 5000 mixtures of normal components and a non-normal tail for each of the 12 simulations as n increases from 1 to 12. GMMchi analysis is carried out on each of the 5000 simulated mixtures for each of the 12 sets of simulations to compare the GMMchi assessment with the known distribution based on the chosen parameters m, std, the number of data points and the proportion of data points in the tail. The accuracy for each n is then the percentage of correct categorizations.

The results of these simulations for a single normal distribution and a non- normal tail are illustrated in Figure 7a. This shows that once the spread of the tail exceeds 2 standard deviations, the accuracy exceeds 90% and plateaus at 97% above 3 standard deviations.

**Figure 7.**
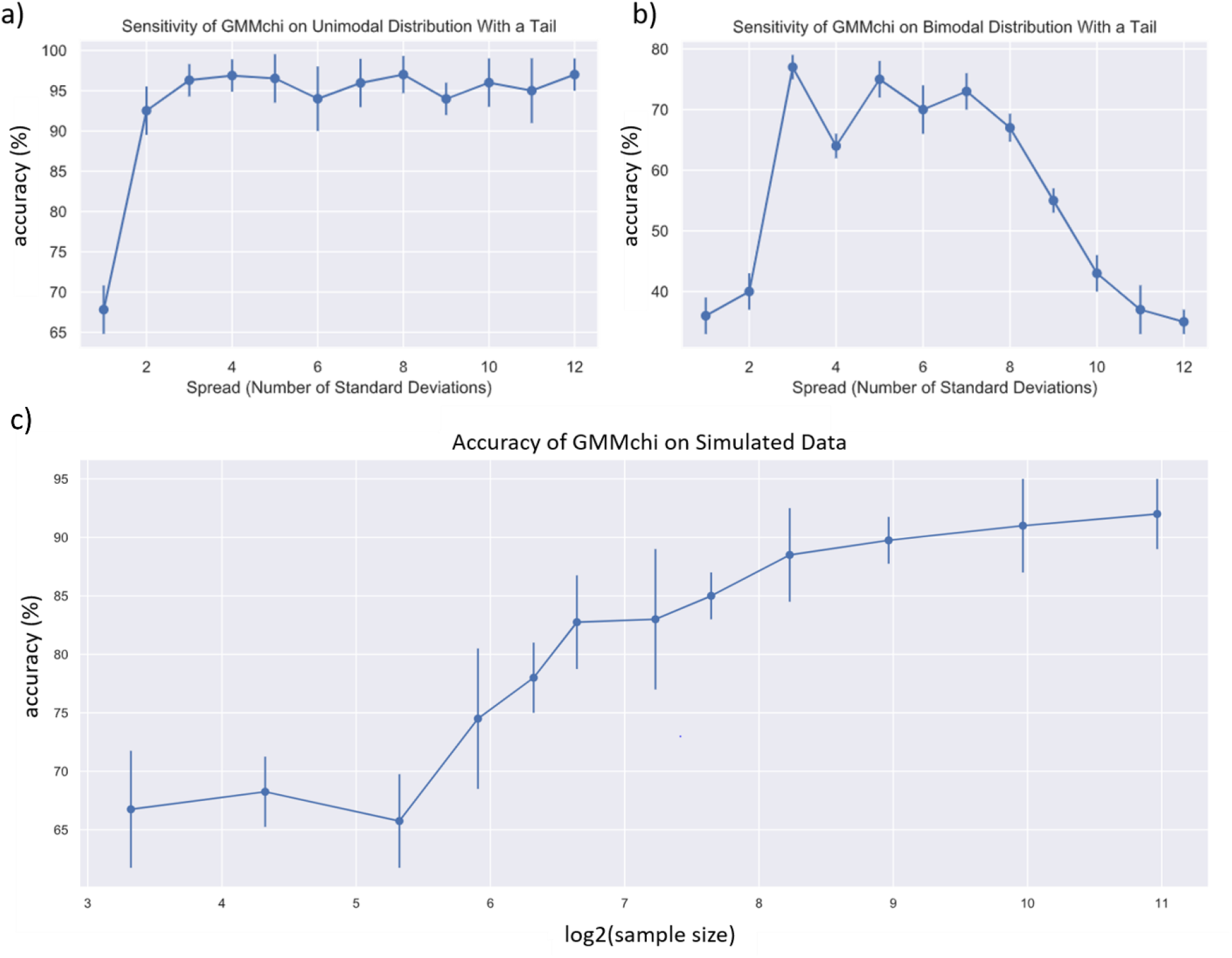
GMMchi Simulated Validation Tests: (a) and (b) show respectively the sensitivity of the GMMchi pipeline for identifying tails with a unimodal distribution or a bimodal distribution. (see the text for details of the simulations and their analysis.) (c) GMMchi’s Accuracy for Categorization of Simulated Distributions for different sample sizes. The graph shows the mean prediction accuracy and its standard deviation in the vertical bars given by the GMMchi pipeline for increasing sample sizes.

For simulated bimodal distributions with a tail Figure 7b shows that the accuracy reaches a plateau of about 70% when the spread of the tail exceeds 2 standard deviations. The accuracy, however, falls rapidly after a spread of 7 standard deviations, suggesting that GMMchi is most sensitive within a tail spread range of 2 to 6 standard deviations. We suspect this is mainly due to GMMchi using a number of bins that is based only the total number of data points. As the tail moves away from the main distribution, in this case, a bimodal distribution, the bimodal distribution will be misrepresented as a unimodal distribution due to the fixed number of bins, leading to the drop in accuracy. We suspect that as the tail spread increases and moves away from the main distribution, the percentage of samples that are predicted by GMMchi to be a unimodal distribution with a tail increase while the percentage of samples that are predicted to be bimodal with a tail drops sharply. This is because with increasing spread of the tail, a high percentage of bimodal distributions with a tail are misclassified as unimodal distributions with a tail. Nonetheless, GMMchi is still able to identify accurately the non-normal tail and delineate the threshold between the main normal components and the tail.

### 4.2 Simulation test 2

The purpose of this simulation is to assess how accurate GMMchi is at predicting the correct categorization with different sample sizes. The four categorizations we included in this simulation are:

1. Unimodal distribution
2. Bimodal distribution
3. Unimodal distribution + tail
4. Bimodal distribution + tail.

We generate a total of 500 distributions, 125 for each of the above four categorizations, using the same basic approach as above for simulation test 1, and do this for 12 different sample sizes. The mean of the normal component(s) is randomly chosen between 3 and 10 and the standard deviation similarly between 0.5 and 5. The number of data points in the tail is randomly chosen to be within 5% to 25% of the total number of data points in the normal component(s). The upper range of the tail is anywhere below 7 standard deviations away from the mean of the upper normal component. After randomly generating 500 simulated mixtures for each of 12 sample sizes from 10 to 2000 we ran GMMchi on each of the mixtures and compared GMMchi’s prediction with the known distribution. The accuracy is, as before, the percentage of correct categorizations. The results of all the simulations are illustrated in Figure 7c. This clearly shows the expected relationship between sample size and average prediction accuracy. GMMchi’s accuracy exceeds 80% with a sample size, n, of about 100 and continues to do well with increasing sample size, exceeding 90% accuracy with a sample size of about 1000. For our microarray data on 78 CRC derived cell lines, which we discuss in the next section, we expect GMMchi to give a prediction accuracy of around 80%.

In the current data landscape, with many datasets having sample sizes well above 100 and often approaching 1000, such as the TCGA CRC data on 637 samples, GMMchi should provide good data models for the objective analysis of observed mRNA expression level distributions.

## 5 Application of GMMchi to Microarray Data from 78 Colorectal Cancer derived Cell Lines

### 5.1 Background Noise Threshold

The background noise threshold is an essential measurement needed when exploring gene mRNA expression data, especially if the data come from a fluorescence-based platform. The background arises necessarily from, for example, the within-array fluorescence noise. An estimate of its level is needed to distinguish convincingly the background from genuine low-level expression. Previous extensive analyses of our microarray data suggested a background noise level between 100 (2^6.65^) and 128 (2^7^). We now use GMMchi analysis of our overall expression data on the 78 CRC derived cell lines to provide an objective statistically based estimate of the background noise level.

The distribution of the microarray expression data for all 54,675 probes on the 78 CRC cell lines is shown in Figure 8a. The number of probes with a given expression level is on the y-axis and the expression levels on the x-axis. The a priori expectation is that this distribution is a mixture of one for probes that are truly -ve on all the cell lines with one for those that express detectable levels of message. The distribution has a shape that looks like a bimodal normal distribution mixing the expected distribution for negative probes with that for positive probes with varying levels of expression, and this is supported by the good fit of a bimodal normal distribution using GMMchi.

**Figure 8.**
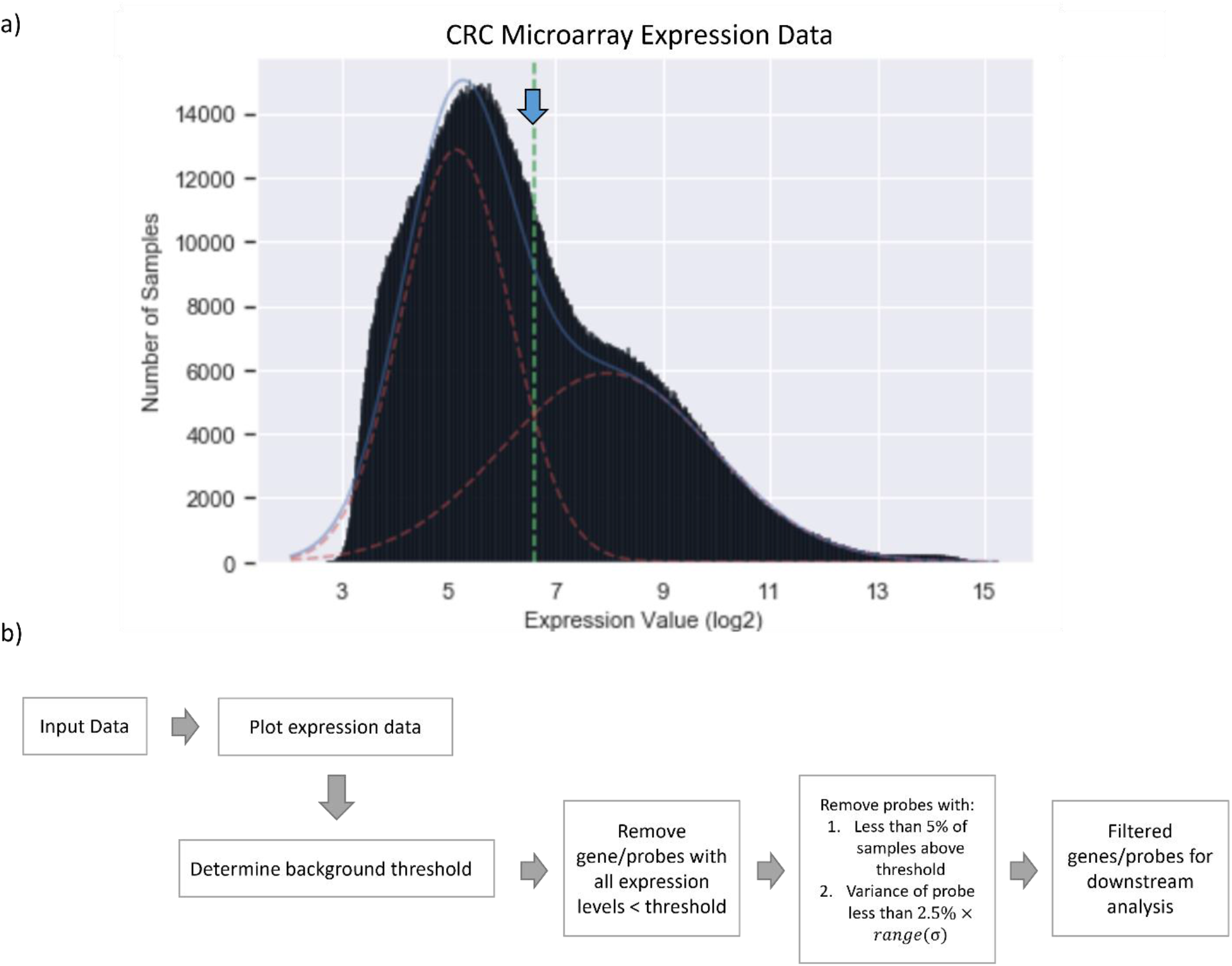
Determining the Background Threshold for Microarray Data Using GMMchi: (a) is the overall distribution of all the expression levels in our microarray data from all the 78 cell lines. The dotted vertical green line and the blue arrow are estimates of the position of the threshold separating the background noise from the true positive expression levels. The estimation is based on a GMMchi fit of two normal distributions to the overall data. (b) is the probe filter pipeline for removing probes and so genes that are predicted to have no significant expression above the background noise threshold.

The intersection between the two normal distributions is used to define the background noise threshold, which on this basis was 2^6.59^ or 96.4, as shown by the vertical green line in Figure 8a. This result agrees with the previous observation of about 2^7^ as a rule of thumb for the background threshold. To clearly distinguish bins expressing background signal from bins with real but low signals, for example in Figure 9, the graphical output of GMMchi colors bins below the threshold 2^6.59^ pink and bins below the threshold 2^7.16^ as yellow. The threshold 2^7.16^ is derived from 2 standard deviations (σ = 1.01) away from the mean (µ = 5.14) of the background distribution to ensure we remove 95% of the background expression level. This ensures more confidence in the identification of genuine positives for further GMMchi analysis.

**Figure 9.**
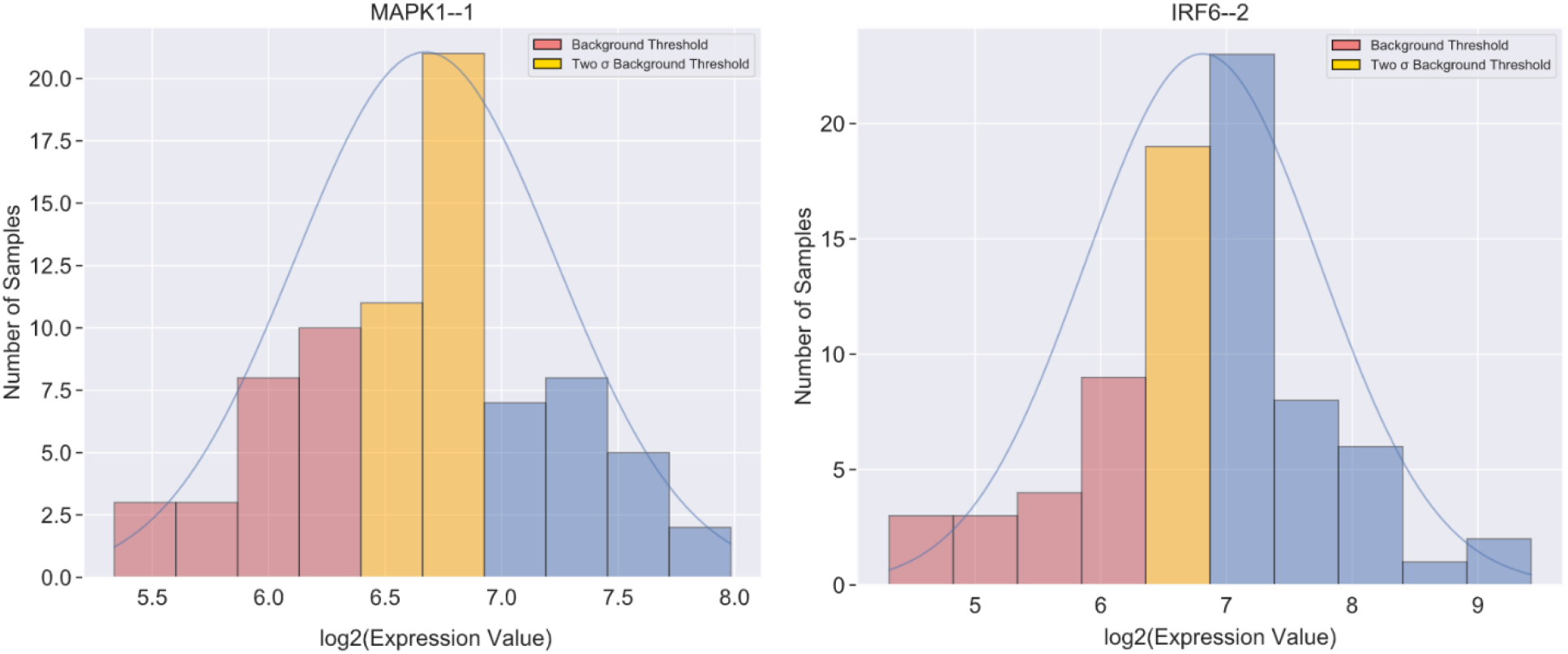
Examples of Genes with different Categories of Unimodal Expression: Both the examples of unimodal distributions have part of their distribution below the threshold for no expression and part above. Pink bins, labeled as ‘Background Threshold’, are below the background threshold (>2^6.59^) while the yellow bins, labeled as ‘Two σ Background threshold’, are samples below 2 standard deviations from the mean of the background distribution estimated from Figure 8a (<2^7.16^).

Therefore, in further analysis of the CRC cell line data we include the background noise threshold to signify lack of expression, namely any expression value less than the background threshold is counted as no expression. This variable is a built-in function in the current GMMchi pipeline. GMMchi analysis thus categorizes cell lines into three groups: background, low, and high, which we aim to associate with different biological functions.

### 5.2 Probe filter

The probe filter removes genes that are considered as non-expressing, following the procedure shown in Figure 8b. Removing genes that are not expressed at all in the cell lines, and so do not contribute to functional interpretation, reduces the computational load.

The function **GMM.probe_filter** first removes genes where all cell lines express levels below the background threshold. The pipeline then searches for genes that are on the borderline of being non-expressing, defined as genes that are expressed above the threshold level in fewer than 5% of the cell lines and whose distribution variance is less than 2.5% of the total range of variances across all genes. These values are arbitrarily chosen as the default values to achieve a stringent criterion for complete lack of expression. They can be easily changed in the algorithm.

### 5.3 Unimodal categorization criterion

We noticed that nearly half of the unimodal distributions had parts of their expression distribution at less than the background threshold, suggesting no expression, and the rest expressed, but with no clear separation between expression and lack of expression (see Figure 9). Inherently, the different groups of genes observed with this pattern of unimodal distribution may also be of interest, though less likely to be associated with clear cut genetic or stable epigenetic changes. While the aim for bimodal distributions is to identify either low versus high expression groups of cell lines or no expression versus clearly expression groups, the expectation for unimodal distributions is to explore whether there is a functionally interesting difference between a non- expressing and a clearly expressing subgroup of cell lines within the unimodal distribution. We call these types of unimodal distributions ‘categorical’. The threshold for categorical unimodal distributions is defined as having less than 95% or more than 5% of the data of a unimodal distribution without a tail above the background threshold of 2^7.16^. The criterion ensures we maximize capturing categorical unimodal distributions while discounting any unimodal distributions that are completely background signal or entirely above background threshold.

### 5.4 Probe Combining Pipeline

In microarray-based expression analysis, the expression level of a gene can be linked to multiple probe sets owing to microarray’s probe-based method of detecting the level of a specific DNA or RNA fragment through groups of probe sets that identify a unique region of a gene (Liu, Bebu, and Li 2010). The probe is then mapped to a gene name using a predefined library consisting of a dictionary of probes to gene translation. Putting this together, each probe set is designed to pick up one message type that aims to identify only one gene based on the message sequence. The data generated by microarray expression analysis where a gene name is associated with multiple probe sets is thus trickier to interpret than RNA-sequencing where a gene name is only associated with a single probe set.

To address and facilitate analysis on a gene-by-gene basis, we developed a probe combining pipeline that aims to combine expression levels of multiple probe sets representing a gene into a single gene with a weighted value that encompasses all the combined probe sets. However, sometimes not all probe sets associated with a given gene can be combined to represent that gene. When we identify probe sets that identify the same gene but do not correlate well with most of the other probe sets, we will retain the probe set without combining it with the rest and assume it is a splice-variant of the gene since we expect splice-variants to behave differently from each other.

As seen in Figure 10a, the pipeline starts with identifying genes with more than one probe set. The pattern of each probe set is predetermined by GMMchi. Bimodal probes consist of 1s and 2s with 1 representing low expressing cell lines and 2 representing high expressing cell lines. In unimodal probes, only probes that exhibit a categorical unimodal pattern are not excluded and consist of 1s and 2s with 1s representing lower expressing cell lines and 2s representing higher expressing cell lines. As mentioned in the last chapter, we are interested in identifying patterns that result in separation into two distinct groups.

**Figure 10.**
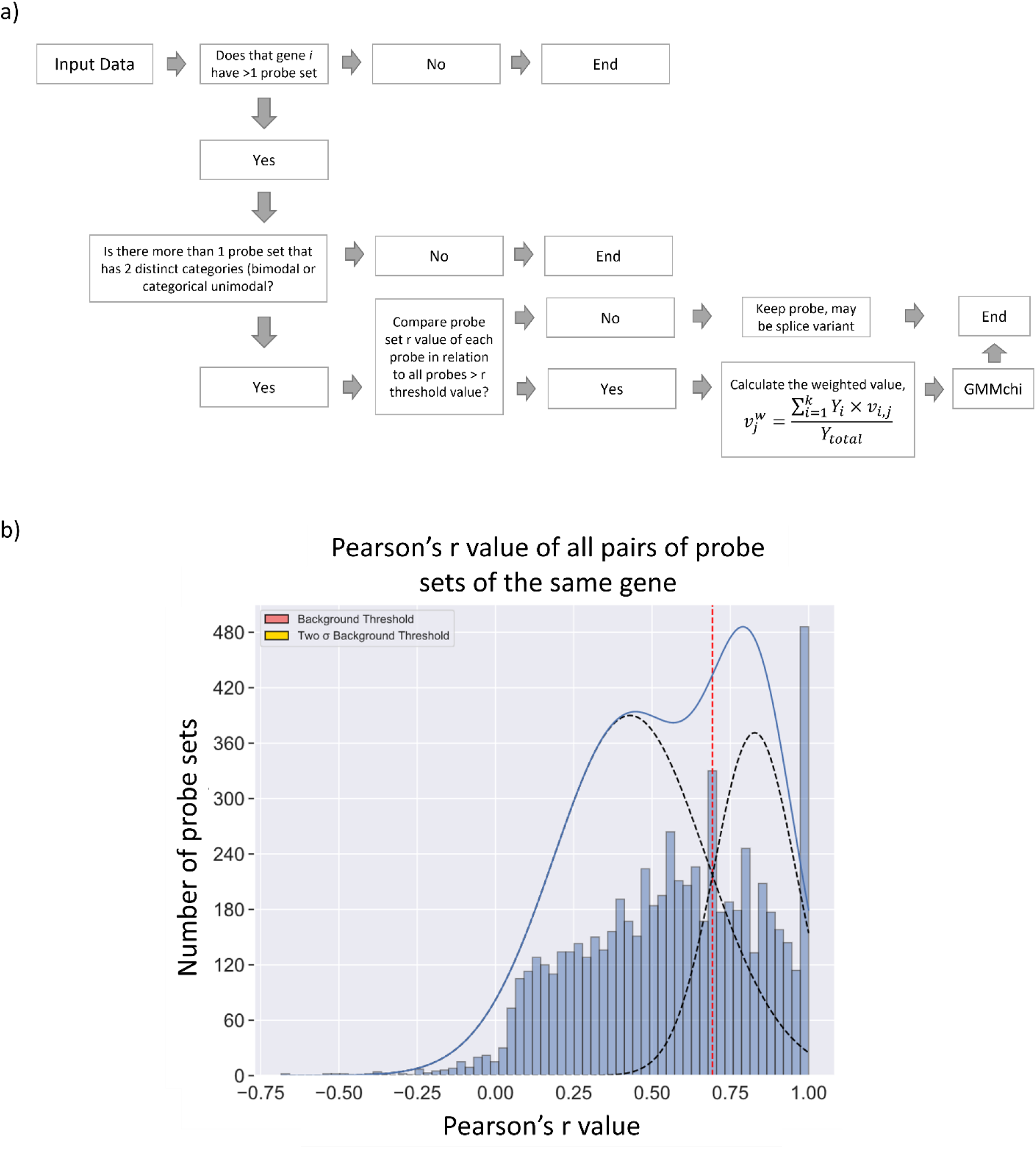
Schematic of the Probe Combining Pipeline: (a) The pipeline for probe combining is used to combine probe sets in microarray data that represent the same gene while keeping together probe sets with a different expression patterns as potential splice variants. (b) shows the total distribution of the r-values for all pairs of probe sets of the same gene. We apply GMMchi to the bimodal distribution which shows the difference between pairs of probes that are associated with each other, at the upper end, and those that are not, the lower distribution. The cutoff that defines the lower r threshold value for probes that are significantly associated with each other is at 0.69, where the two fitted distributions intersect, as indicated by the vertical red dotted line.

The association between different probe sets from the same gene is determined by Pearson’s r correlation. A group of associated probe sets is identified as all those with pairwise correlations above 0.69, the threshold that indicates a significant association determined by the cutoff of the bimodal distribution identified by plotting out all the r-values using GMMchi, as shown in Figure 10b. A weighted mean for the group is calculated using as weights the inverse of corresponding chi-square goodness of fit value returned from the GMMchi output for each probe set member of the group. Thus if

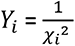 is the inverse of the chi-square goodness of fit value from GMMchi for probe *i*

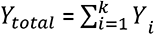 is the sum of all inverse chi-square goodness of fit values for the k probe sets belonging to the group, and

*ν_i,j_* expression value of probe *i* for a given cell line *j,* then

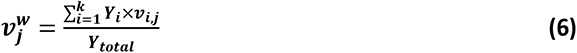

is the weighted expression level for this group of probe sets for line j. The inverse of the goodness of fit value *Y_i_* is used as the weight for each probe *i* expression value because the lower the goodness of fit value the better the fit. If all the weights are the same, then the weighted mean is just the arithmetic mean of the k expression values. The weighted mean is then put through GMMchi to convert those values into groups of 1 (low expression) or 2 (high expression) for further analysis.

Lastly, probe sets that do not behave similarly with other probe sets (r-value < r threshold value) are treated as separate splice variants and kept as a separate probe set annotated with the gene plus the probe number (i.e., *CDH1--2*). The result of this pipeline is then the basis for further downstream analysis.

### 5.5 Examples of specific analyses

Figure 11 shows two examples illustrating the power of GMMchi for categorizing continuous microarray gene expression data. The distribution for gene *EPHB3*, which encodes Ephrin Receptor B3, a receptor tyrosine kinase involved in mitogenic and differentiation signaling, is shown in Figure 11a. A clear cutoff at 9.4 separates the mixture of two normal components. The Q-Q plot supports the assumption that the two component distributions are indeed normal. The expression distribution for the gene *CDH1*, which encodes E-Cadherin, a key epithelial cell integral membrane protein, is shown in the first histogram in Figure 11b. The initial GMMchi fit shows a normal distribution including a majority of the 78 cell lines expressing high levels of E-Cadherin with a second distribution for a small group of lines expressing low or no levels. The application of the tail criterion suggests that the small low-level group should be considered as a non-normal tail, as shown by the second and third histograms in Figure 11b. The Q-Q plot in the final panel of Figure 11b then confirms the final result of GMMchi as a normal distribution of positively expressing cell lines with a non-normal tail of low or non-expressing lines.

**Figure 11.**
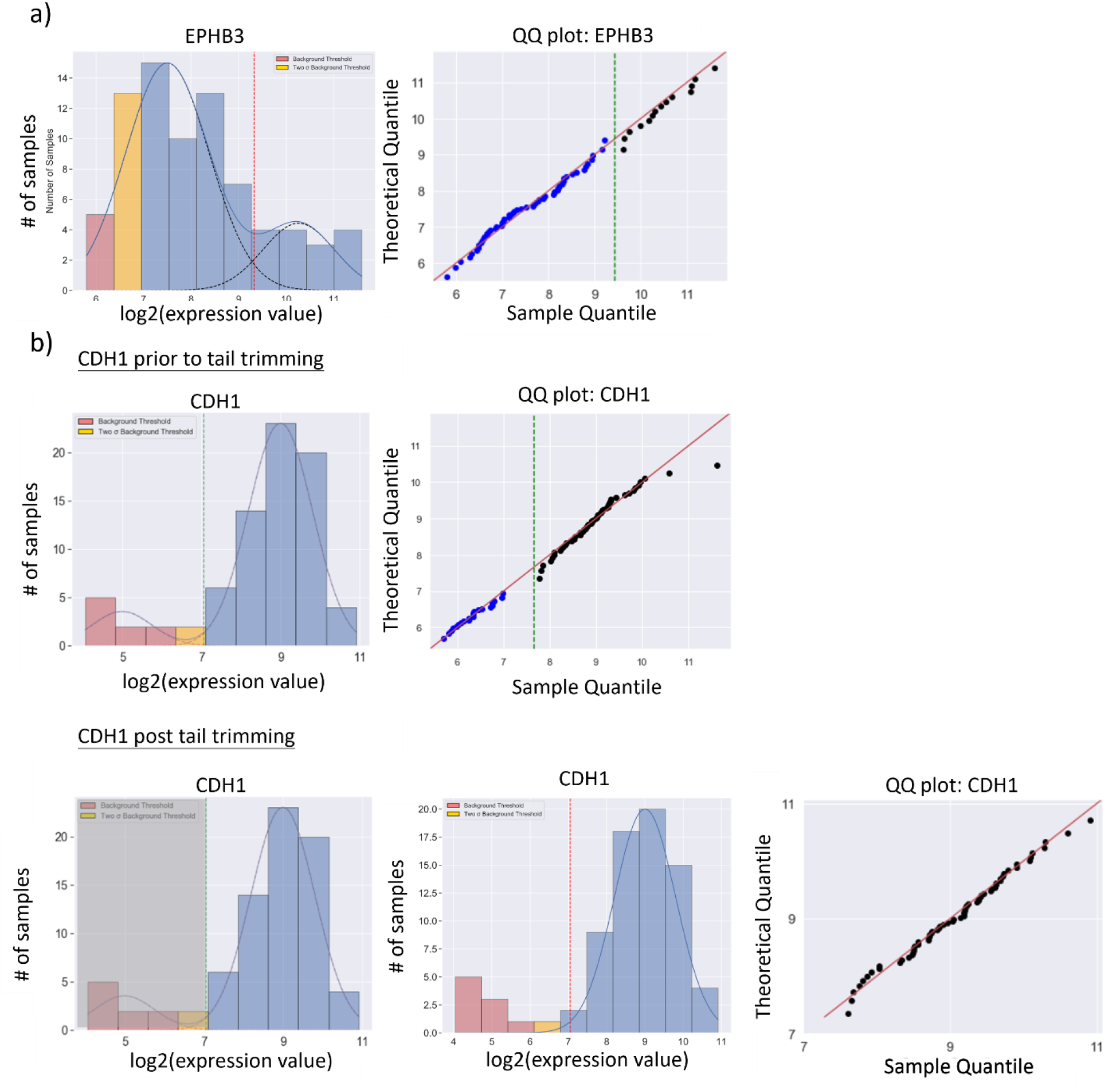
GMMchi on Gene Expression Data: As before, in both cases, the pink columns indicate clearly negative expression while the yellow columns indicate at most very low-level expression. The dotted red lines mark the intersection between the two fitted normal curves (a) or between the fitted single normal distribution and the tail (b), and so the best estimate of where to separate low probably negative expressing cell lines from those that are clearly positive. The expression distribution in (a) is for gene EPHB3 encoding the protein Ephrin Receptor B3. In this example, GMMchi estimates the cutoff between the mixture of two normal components that are adequately normal as shown in the Q-Q plot with the data along the 45-degree line. In (b), a good example of a tail trimming process, the first histogram shows the expression distribution for CDH1, the epithelial membrane protein E-Cadherin, prior to tail trimming. The first Q-Q plot revealed datapoints deviating from the 45-degree line, suggesting an inadequate fit. The second histogram shows the tail identification step where the grey shading indicates the potential identification of a non-normal tail. The third histogram is the result of iterative tail trimming while the last figure shows the Q-Q plot of the resulting fit, indicating an adequately fitted normal component with a non-normal tail.

### 5.6 Lumen-forming CRC Cell Lines and CDX1 associations

A colorectal cancer can be a mixture of cell types consisting of different stages of differentiation. Even though tumors are known to have a higher percentage of cells that are the stem cell population, there can also be different subpopulations of differentiated cell types. These can be any of the three major differentiated colonic cell types, enterocytes, goblet cells, and enteroendocrine cells. Here, we explore genes associated with lumen formation and enterocyte differentiation.

It was previously shown that caudal-related protein 1 (*CDX1*) is the primary transcription factor responsible for enterocyte differentiation with, however, a strong association with the closely related *CDX2* (Ashley et al. 2013). Figure 12 shows the GMMchi analysis for the expression of *CDX1* and *CDX2,* and three other differentiation markers, carcinoembryonic antigen-related cell adhesion molecule 5 (*CEACAM5,* the original *CEA*), villin 1 (*VIL1*), and glycoprotein A33 (*GPA33*). All these 5 gene expressions have clear bimodal distributions making them good candidates for 2x2 association analysis using the function GMM.run_hits. The 2x2 Table 1a shows the close association between the expression of *CDX1* and the 4 other genes. Note that the association between the expressions of *CDX2* and *CDX1* is an inclusion, namely with all *CDX1* high being *CDX2* high, but not the reverse.

**Figure 12.**
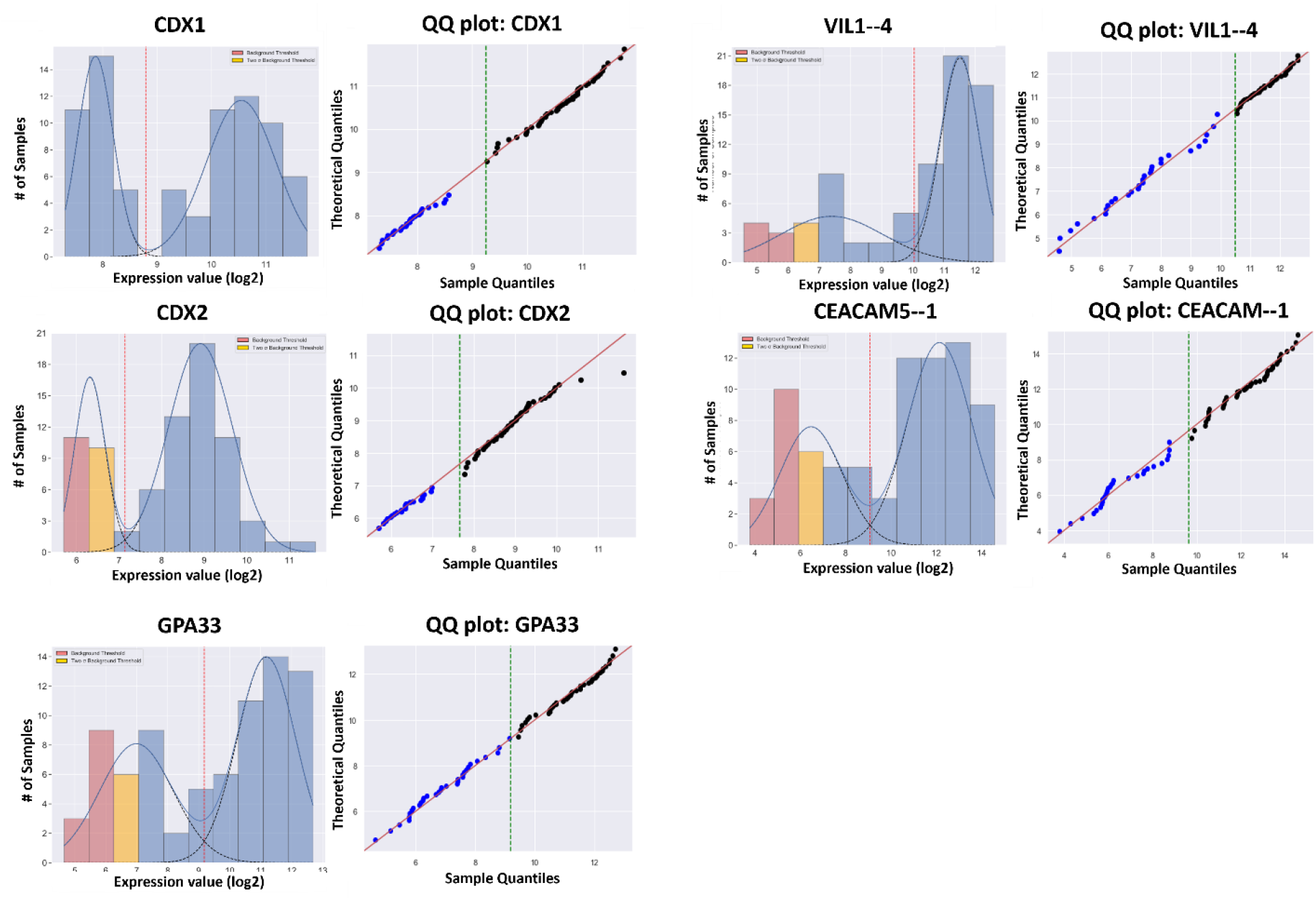
Examples of GMMchi Analysis on Known Differentiation Markers: Here we show examples of known differentiation markers that are positively and significantly associated with CDX1, a biomarker for colonic epithelial differentiation identified by phenotype association with lumen formation. All genes exhibited a well-defined bimodal distribution of expression levels consisting of two subpopulations.

**Table 1.** Genes significantly associated with CDX1 and lumen formation: The R-values from 2x2 association analyses between the genes whose expression is illustrated in Figure 12. The p- values from the χ^2^s for each pairwise comparison are all Benjamini–Hochberg corrected. (a) Shows that CDX1 expression is strongly correlated with those of CDX2 and GPA33, a basolateral intestinal differentiation marker, as well as with CEACAM5 and VIL1 which are also colonic epithelial differentiation markers. The association between CDX1 and CDX2 is an inclusion, namely with all CDX1 high being CDX2 high, but not the reverse. (b) Shows the 2x2 analyses for the 4 gene expressions that are most highly correlated with lumen formation. Lumen formation is included within CDX1and CDX2 high expression, while NRCAM-1 included within lumen formation

| **CDX1** | **+/+** | **+/-** | **-/+** | **-/-** | **p-value** | **R value** |
| --- | --- | --- | --- | --- | --- | --- |
| GPA33 | 43 | 4 | 3 | 28 | 1.03E-13 | 0.81 |
| CDX2 | 47 | 0 | 8 | 23 | 2.29E-13 | 0.80 |
| CEACAM5—1 | 43 | 4 | 6 | 25 | 6.37E-11 | 0.73 |
| VIL1—4 | 42 | 5 | 7 | 24 | 1.98E-09 | 0.68 |

b)
| **Lumen Formation** | **+/+** | **+/-** | **-/+** | **-/-** | **p-value** | **R value** |
| --- | --- | --- | --- | --- | --- | --- |
| NRCAM—1 | 12 | 10 | 1 | 19 | 6.38E-04 | 0.53 |
| CDX1 | 19 | 3 | 9 | 11 | 7.99E-03 | 0.44 |
| CDX2 | 20 | 2 | 11 | 9 | 1.32E-02 | 0.42 |
| CEACAM5—1 | 18 | 4 | 9 | 11 | 2.23E-02 | 0.38 |

In total there are 22 out of 42 cell lines with the phenotype lumen formation as assessed from previous experiments by lumen formation assays (Ashley, Yeung, and Bodmer 2013). Table 1b shows the 2x2 analyses for the 4 gene expressions that are most highly correlated with lumen formation. As expected, this includes the genes *CDX1, CDX2* and *CEACAM5,* all as ’including’ lumen formation in the sense that all the lumen forming lines expressed these three genes at the high level but not the reverse. The expression of the *CDX1* associated genes *GPA33* or *VIL1* was not found to be correlated with lumen formation at this high level of significance. However, the expression of the neuron-glia-related cell adhesion molecule (*NRCAM*) was significantly associated with lumen formation as an inclusion. It seems likely *NRCAM*, a differentiation marker for neural development, is a nonessential gene for lumen formation that can nevertheless be expressed in lumen-forming cells as a sort of bystander effect (Conacci-Sorrell et al. 2002).

These analyses show how associations between continuous data, such as the gene expression data, can be related to categorical data such as, for example, mutations, distinctive phenotypes, and methylation, by transforming the continuous data into categorical data using GMMchi and so revealing interesting patterns of gene expressions associated with patterns of differentiation.

### 5.7 MUC2-expressing Goblet Cell Subpopulation in CRC Cell Lines

Goblet cells are one of the three differentiated cell types in the colon and are responsible for the production of mucin 2, MUC2, the major mucin that lines the intestine. The GMMchi analysis of *MUC2* (the gene for MUC2) and three other genes (*TFF3, REG4* and *FCGBP*) known to be involved in normal goblet cell differentiation are shown in Figure 13. The QQ plots for GMMchi for *TFF3, REG4,* and *FCGBP* in Figure 13 show multiple data points deviating from the 45-degree line near the cutoff threshold (green dashed line). This is mainly because of the relatively high overlap between the two normal distributions, making the membership of the data points within the overlap harder to assign and thus resulting in a relatively poor separation between the two distributions. This may be due to low expression of these three genes being due to a small proportion of expressing cells in a cell line rather than an overall low level of expression on all the cells. TFF3 and FCGBP are proteins that seem to act as chaperones for the maturation of the mucus and our evidence suggests the existence of a significant subset of CRCs that express TFF3 and other proteins associated with mucus maturation, but not the mature MUC2 itself. The biology of this is now being further investigated.

**Figure 13.**
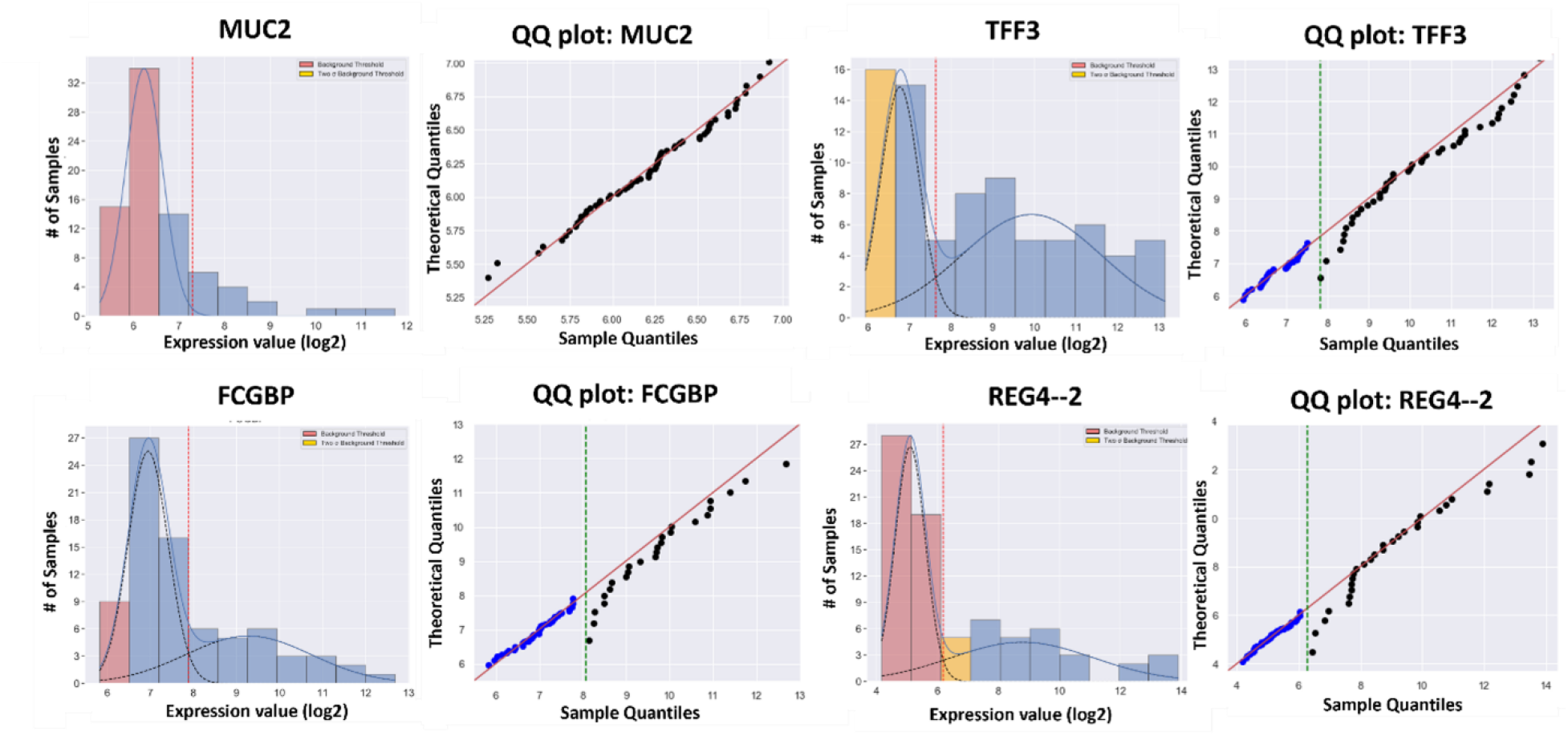
GMMchi Analysis on Genes Associated with Goblet Cells: (a) GMMchi analysis of *MUC2* and three other genes (*TTF3, REG4* and *FCGBP*) known to be associated with goblet cell differentiation.

The GMMchi pipeline allows the possibility of identifying given cell lines or subsets of the cell lines in the histogram output. Two in-depth examples of this are given in Figure 14 and show how such analysis can reveal novel subsets of the cancers which may be of interest for further experimental investigation. The subset identification in Fig 14a very clearly shows that almost all MUC2+ are TFF3+ while there exists a major subset of lines with TFF3+ and no MUC2, as already noted. The two cell lines (shown in green) that are apparently MUC2+TFF3- turn out, by immunofluorescence (unpublished data), to have very low proportions of TFF3+ cells, emphasising the problem of detecting such low levels of expression by mRNA analysis of the bulk cell line preparations. Signet-ring cancers are a rare subset of CRCs with very high MUC2 and TFF3 expression. Figure 14b shows that almost all MUC2 expressing cell lines also express high levels of FCGBP, whilst a significant proportion of the TFF3+ cell lines express low or no FCGBP.

**Figure 14.**
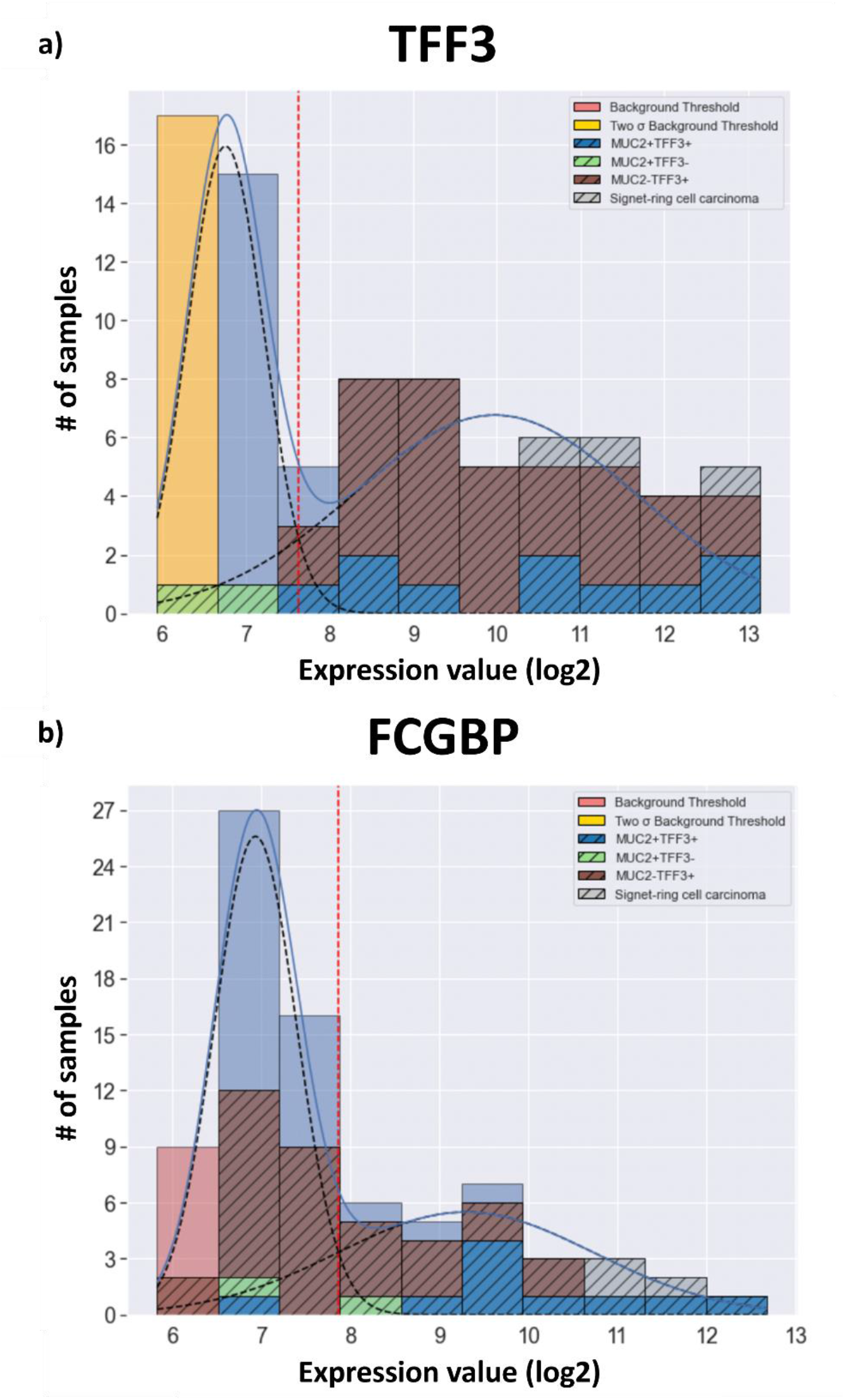
Examples of GMMchi outputs with defined subsets of cell lines: (a) TFF3 expression analysis with defined subsets, MUC2+TFF3+, MUC2+TFF3-, MUC2-TFF3+ and Signet-ring carcinomas, shown by the indicated coloured crosshatched segments. (b) FCGBP expression analysis with with defined subsets, MUC2+TFF3+, MUC2+TFF3-, MUC2-TFF3+ and Signet-ring carcinomas, shown by the indicated coloured crosshatched segments.

In Table 2, the 2x2 analyses of top 50 genes significantly associated with *MUC2* expression are obtained using the function **GMM.crosstab_table**. Several genes whose function has been shown or suggested to be associated with goblet cell differentiation, and so MUC2 production, occur in this list They include, for example, REG4, a secreted protein found to associate closely with secretory cells as well as playing an important role in intestinal morphogenesis (Sasaki et al. 2016), *FCGBP*, Fc- γ binding protein suggested to crosslink with MUC2 and have a chaperone like activity on the maturation of MUC2, *ST6GALNAC1 and UGT1A8 /// UGT1A9*, involved in glycosylation of MUC2 (Bergstrom and Xia 2013), *AGR3,* whose product is involved in catalyzing disulfide bond formation in protein folding and so contributes to maintaining the structure of MUC2, *CEACAM5,* associated with both goblet cell and enterocyte differentiation, and the transcription factor *ETS2*. A trefoil factor TFF1 is also in the list, and we have included the trefoil factor TFF3, though it comes just below the cutoff point for the top 50 associations, because of its well known almost total association with MUC2 expression in normal colonic tissue. The trefoil factors are also known to play an important role in MUC2 maturation, possibly after its secretion (Kim and Ho 2010). Most of the associations are inclusions suggesting functions that may be required for goblet cell differentiation and MUC2 maturation but that also have other roles. Further experimentation is being carried out to explore the potential role of other genes in the Table 2 list in goblet cell differentiation and MUC2 maturation, and to elucidate the significance of the relatively large proportion of CRCs that seem to be on their way to goblet cell maturation by the expression of TFF3 but do not produce mature MUC2.

**Table 2.**
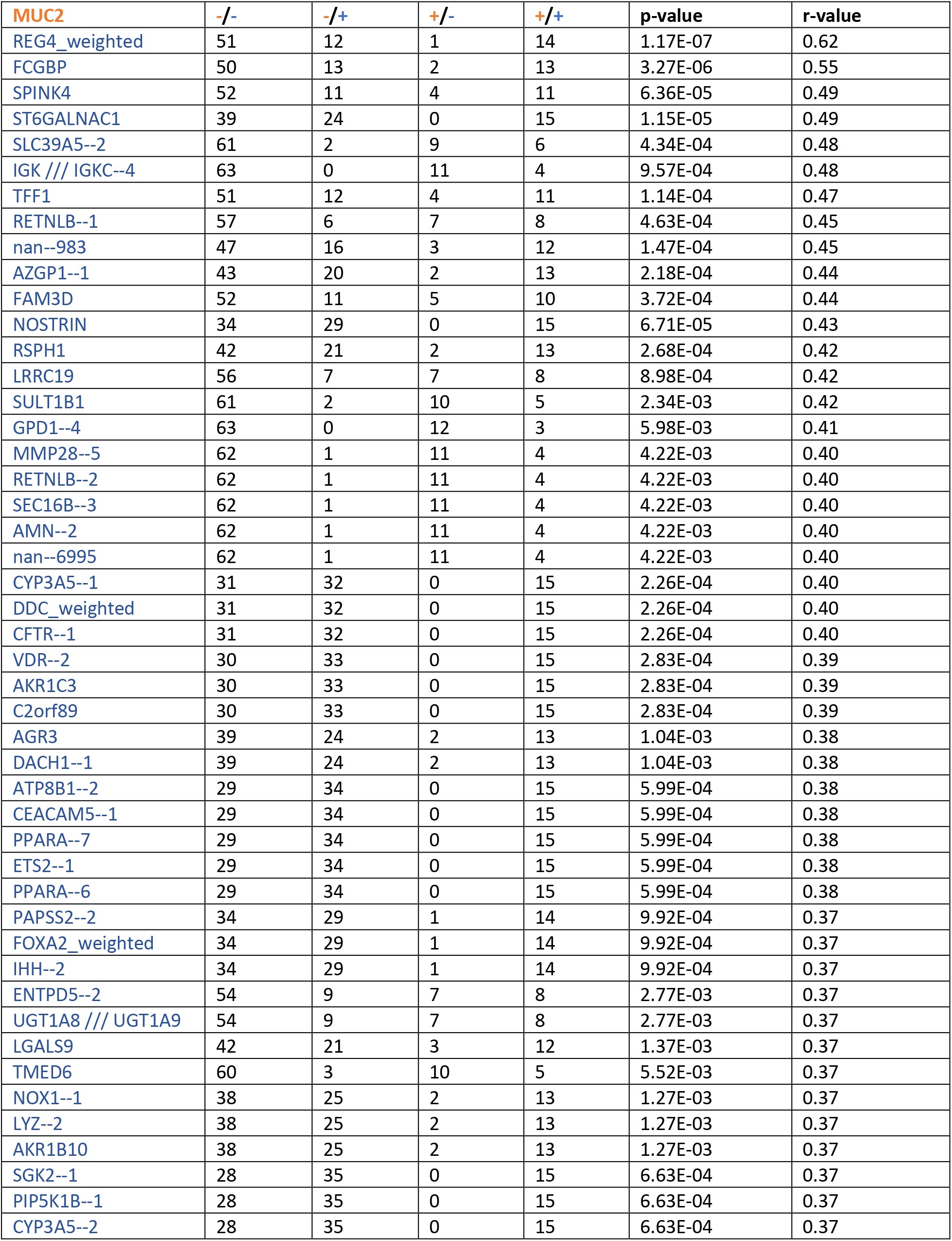
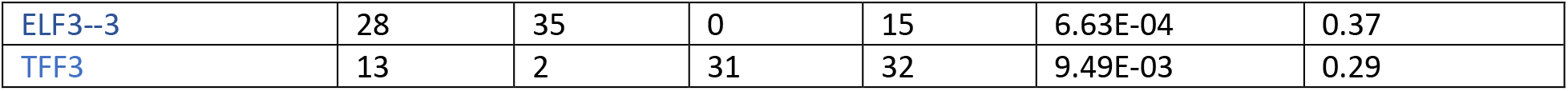
Top 50 Genes Significantly Associated with MUC2: Genes positively correlated with MUC2 in 78 colorectal cancer cell lines.

### 5.8 Methods summary

A python library was built for easy utilization of the GMMchi pipeline. Built-in methods (functions) within the pipeline include:

a. **GMM.GMM_modeling**: This is used to perform GMMchi on an input matrix of samples x features. The input is a continuous mRNA measurement using either RNA-Seq or microarray (as discussed in the main text).
b. **GMM.probe_filter**: This is used to perform probe filtering to remove genes that are considered not to be expressed (as discussed in the main text).
c. **GMM.find_hits**: This is used to calculate 2x2 contingency tables from the categorized data returned from the GMM.GMM modeling output.
d. **GMM.run_hits**: This is used to output the full 2x2 contingency table from a pre- defined set of genes of interest (as discussed in the main text).
e. **GMM.crosstab_table**: This is used to visualize the full 2x2 contingency table from a pre-defined set of genes of interest (as discussed in the main text).

The whole program is implemented in Python and can be imported as a library; the full code is shown in the appendix, the library and installation link are online at: https://github.com/jeffliu6068/GMM

## 6 DISCUSSION

The somatic mutations or stable epigenetic changes whose selection drives cancer progression are discrete genetic variations analogous to the germ line genetic variants that are inherited following Mendelian segregation. The consequences of these somatic genetic changes can be seen in changes in the levels of mRNA expression of a wide range of genes. These levels are continuous variables. At the simplest level there could, for example, be a significant change in the message level of a mutated or epigenetically altered gene. A collection of cancers, some of which carry this genetic change and some not, would then be expected to have a bimodal distribution of the message levels for this gene, giving rise to a dichotomised phenotype of high versus low levels of expression. The more closely associated the function of the altered gene, for example having a role in the control of a particular cellular differentiation process, is with other genes also involved in the same function, the more likely it is that these other genes will have similar correlated bimodal distributions to that observed for the altered gene.

The aim of this paper has been to create a widely applicable and flexible tool to analyse the relationship between discrete genetic changes and continuously variable gene expression levels by transforming the continuous expression levels into dichotomised phenotypes. Intrinsically dichotomous phenotypes, such as presence or absence of a particular cell type or ability to form a particular differentiated structure or not, can then be included with the derived dichotomised gene expression phenotypes for further analyses of their interrelationships.

The obvious first step is to apply GMM (Gaussian Mixture Modelling) to observed gene expression levels for a range of cancers, assuming, as has been widely observed that, when log_2_ transformed, these usually have a distribution that is well described by a normal distribution. When, however, as is likely for cancers a relatively small subset of the cancers has either a higher or lower expression level than their normal counterparts, these distributions are less likely to be normal. This is the ’tail’ problem, and so a primary goal of the analysis has been to find an objective procedure for identifying non-normally distributed tails. This has been done by a combination of iterative tail pruning and GMM, using a χ^2^ goodness of fit approach to enable the selection of the best fitting mixture of a GMM derived distribution and a non- normal tail. The success of this approach, GMMchi, has been demonstrated using genome wide microarray derived gene expression data from a collection of 78 CRC (colorectal cancer) cell lines. The complete pipeline starts by filtering out genes that are not expressed in any cell line, or those that are well described by a single unimodal normal distribution and allows for detection of splice variants when multiple oligonucleotide sets are used for a single gene. GMMchi can be applied to obtain dichotomised phenotypes from any similar source of data, such as single cell mRNA data or TCGA RNAseq data, whose analysis will be the subject of a further paper.

Simulated mixtures of normal distributions and tails confirm the overall consistency of our pipeline, indicating it reaches an accuracy of 85% once the number of samples is sufficiently large at around a minimum of 90 samples, which is close to the number of cell lines in our demonstration data set. The accuracy of identifying a bimodal distribution with a tail spread beyond 7 standard deviations decreases quite rapidly with increasing tail size. It is, however, rare in practice to find tails that span beyond 7 standard deviations from the main component in gene expression data.

The GMMchi pipeline iterative tail pruning process so far allows for only a single tail at either the upper or lower end of the overall distribution. This could readily be extended to allow tails at both the upper and lower ends, though we did not see evidence for the need for this in our analysis of the CRC cell line data.

The 2x2 analyses associated with the analysis of the lumen formation and goblet cell differentiation phenotypes can be extended to searches for clusters of gene expressions not so far clearly associated with any defined phenotype. If there are different types of continuous measurements on the same set of cell lines, for example mRNA by microarray analysis and RNAseq, then each data set can be separately dichotomised by GMMchi giving rise to discrete phenotypes that can also be jointly analysed. Missing data are simply accounted for in a 2x2 analysis by only including pairs of cell lines that have data for both pairs of dichotomised phenotypes. Thus, for all those genes with at least two defined levels of expression, which if there are more than two levels can be condensed into just two levels, it is possible to look for associated clusters using 2x2 analyses with a defined minimum high-level r or low-level p value cut off, using the r values as a distance measure for clustering. A cluster identified in this way can be turned into a further dichotomised phenotype by assigning a cell line to a high or low category depending on the consistency of its expression of the members of the cluster at either a high or a low level. Such phenotypes could themselves then be used for further 2x2 analysis. This is an approach used for the early identification of HLA types using antisera from multiparous women known, mostly, to have antibodies to two or more HLA specificities (Payne et al., 1964).

In conclusion, we propose that GMMchi is a general tool for the analysis of the interrelationships between a matrix of measurements of a set of continuous variables on a panel of objects, based on dichotomisation of the continuous variables and the analysis of their 2x2 associations.

